# Functional clusters of neurons in layer 6 of macaque V1

**DOI:** 10.1101/685990

**Authors:** Michael J Hawken, Robert M Shapley, Anita A Disney, Virginia Garcia-Marin, JA Henrie, Christopher A Henry, Elizabeth M Johnson, Siddartha Joshi, Jenna G Kelly, Dario L Ringach, Dajun Xing

**Affiliations:** Center for Neural Science, 4 Washington Place, New York University, NY, NY 10003; Dept. Neurobiology, Duke University, Durham, North Carolina 27110; Dept. Neuroscience, Albert Einstein College of Medicine, Bronx, New York 10461; Wharton Neuroscience Initiative, University of Pennsylvania, Philadelphia, PA 19104; Dept. Neuroscience, University of Pennsylvania, Philadelphia, PA 19104; Dept. Neurobiology, David Geffen School of Medicine at UCLA, Los Angeles 90095; Key State Laboratory of Cognitive Neuroscience and Learning, Beijing Normal University, Beijing 100875

## Abstract

Layer 6 appears to perform a very important role in the function of macaque primary visual cortex, V1, but not enough is understood about the functional characteristics of neurons in the layer 6 population. It is unclear to what extent the population is homogeneous with respect to their visual properties or if one can identify distinct sub-populations. Here we performed a cluster analysis based on measurements of the responses of single neurons in layer 6 of primary visual cortex to achromatic grating stimuli that varied in orientation, direction of motion, spatial and temporal frequency, and contrast. The visual stimuli were presented in a stimulus window that was also varied in size. Using the responses to parametric variation in these stimulus variables we extracted a number of tuning response measures and used them in the cluster analysis. Six main clusters emerged along with some smaller clusters. Additionally we asked whether parameter distributions from each of the clusters were statistically different. There were clear separations of parameters between some of the clusters, particularly for f1/f0 ratio, direction selectivity, and temporal frequency bandwidth but other dimensions also showed differences between clusters. Our data suggest that in layer 6, across the spatial extent of a single cortical hypercolumn, there are multiple parallel circuits that provide information about different aspects of the visual stimulus.

## Introduction

The study was motivated by the growing realization that layer 6 has multiple functional roles in the visual function of macaque primary visual cortex, V1. Layer 6 provides major output from V1 to extrastriate (Fries et al, 1985) and subcortical targets (Fitzpatrick et al, 1994) that have different functional characteristics. Layer 6 also has multiple distinct intracortical recurrent pathways and provides long range connections within the infragranular layers of V1. Layer 6 feedback afferents (McGuire et al, 1984; Wiser & Callaway, 1996) provide a considerable fraction of the excitatory synapses in V1 layer 4C. Furthermore, such layer 6 feedback to layer 4C is provided by distinct subpopulations (Wiser and Callaway, 1996) and appears to be of considerable importance in the generation of the dynamical responses within layer 4Cα (Chariker et al, 2016; 2018).

The goal of the current study was to examine layer 6 neuron responses along multiple stimulus dimensions to seek functional clusters of layer 6 neurons by means of hierarchical clustering analysis. As shown in Results, distinct functional clusters emerged in the analysis. After presenting the functional clusters, we discuss their possible functional roles and the relation of functional clusters to classes of layer 6 neurons with different anatomical circuit motifs (Wiser and Callaway, 1996; Briggs and Callaway, 2001).

Although layer 6 in the primate has a number of distinct morphological neuronal populations (Callaway, 1998; Briggs et al, 2016), previously it was concluded that there is little information that suggests there are distinct functional populations (cf. Briggs, 2010; Thomson 2010). However, there are indications of functional subpopulations in the anatomical data. Most layer 6 excitatory neurons have intracortical axonal projections (Lund and Boothe, 1975; Fitzpatrick et al, 1985; Wiser and Callaway, 1996) and some of those intracortically projecting neurons have M- or P-pathway-specific axonal projections – targeting either layer 4cα or 4cβ and 4a. There are other layer 6 sub-populations that project specifically to extrastriate areas including MT (Lund et al, 1975; Fries et al, 1985; Nhan and Callaway, 2012). Furthermore, the population of neurons that provide feedback to the LGN (Hendrickson et al, 1978) – about 15% of the neurons in layer 6 (Fitzpatrick et al, 1994) – could be divided into three distinct clusters based on their conduction latency, their maximum response rate, and whether they were simple or complex cells (Briggs and Usrey 2009; Hasse and Briggs, 2017). Other earlier findings also indicate the possibility of functional sub-groups within the layer 6 population: 1) layer 6 neurons are more likely to be direction-selective than the overall V1 population (Livingstone and Hubel, 1984; Orban et al., 1986; Hawken et al, 1988), 2) they show weak extra-classical receptive field suppression (Gilbert, 1977; Schiller et al, 1976a; Sceniak et al, 2001; Henry et al, 2013), 3) there are distinct populations of layer 6 simple and complex cells (Schiller et al, 1976a; Ringach et al, 2002) and 4) there is a range of contrast sensitivity functions in the layer 6 population indicative of M and P cell pathway dominance (Hawken et al, 1996).

In Results, we report that six main functional clusters emerged in the analysis of functional properties of the layer 6 population. The cluster analysis employed seven tuning and response measures: 1) orientation bandwidth (obw), 2) direction selectivity (dI), 3) spatial frequency bandwidth (sfbw), 4) temporal frequency bandwidth (tfbw), 5) contrast threshold (cTh), 6) the contrast where the response reached 50% of the maximum (c50), and 7) modulation index (f1/f0 ratio). The results imply that V1 cortex has very distinct functional classes of neurons within layer 6, a result that indicates there are multiple parallel circuits within a single cortical hypercolumn, each of which provides information about different features of the visual stimulus.

## Methods

### Preparation

Adult male old-world monkeys (*M. fascicularis*) were used in acute experiments. The animal preparation and recording were performed as described in detail previously (Hawken et al. 1996; Ringach et al. 2002; Xing et al. 2005; Henry and Hawken, 2013). Anesthesia was initially induced using ketamine (5-20 mg/kg, i.m.) and was maintained with isofluorane (1-3%) during venous cannulation and intubation. For the remainder of the surgery and recording, anesthesia maintained with sufentanil citrate (6-18 µg/kg/h, i.v.). After surgery was completed, muscle paralysis was induced and maintained with vecuronium bromide (Norcuron, 0.1 mg/kg/h, i.v.) and anesthetic state was assessed by continuously monitoring the animals’ heart rate, EKG, blood pressure, expired CO_2_, and EEG.

After the completion of each electrode penetration, 3–5 small electrolytic lesions (3 μA for 3 s) were made at separate locations along the electrode track. At the end of the experiments, the animals were deeply anesthetized with sodium pentobarbital (60 mg/kg, i.v.) and transcardially exsanguinated with heparinized lactated Ringer’s solution, followed by 4 L of chilled fresh 4% paraformaldehyde in 0.1 M phosphate buffer, pH 7.4. The electrolytic lesions were located in the fixed tissue and electrodetracks were reconstructed to assign the recorded neurons to cortical layers as described previously (Hawken et al., 1988).

All experimental procedures were approved by the New York University Animal Welfare Committee and were conducted in compliance with guidelines in National Institutes of Health’s *Guide for the Care and Use of Laboratory Animals.*

### Characterization of visual properties of V1 Neurons

We recorded extracellularly from single units in V1 using glass-coated tungsten microelectrodes. Action potentials were discriminated and recorded as described in Henry and Hawken (2013). Each single neuron was stimulated monocularly through the dominant eye (with the non-dominant eye occluded). The steady-state response to drifting gratings was determined to provide characterization of visual response properties. For all neurons included in this study measurements of orientation tuning, spatial and temporal frequency tuning, contrast response with achromatic gratings, and area summation curves were obtained. Receptive fields were located at eccentricities between 1 and 6 deg. Stimuli were presented using custom software on either a Sony Trinitron GDM-F520 CRT monitor (mean luminance 90-100 cd/m^2^) or an Iiyama HM204DT-A CRT monitor (mean luminance 60 cd/m^2^). The monitors’ luminance was calibrated using a spectroradiometer (Photo Research PR-650) and linearized via a lookup table using custom software. Stimuli were displayed at a screen resolution of 1024×768 pixels, a refresh rate of 100 Hz, and a viewing distance of 115 cm.

For all tuning measurements the responses were recorded to drifting sinusoidal gratings. In the current study we used the f0 response for neurons with f1/f0 ratios < 1 and the f1 response for neurons with f1/f0 ratios > 1. All fitting was done using the Matlab function fmincon (Matlab) where the least-squared error was used to minimize the objective function.

#### Orientation tuning and direction selectivity

The responses of each neuron were recorded to different orientations between 0 – 360 deg., either in 20 or 15 deg steps. The stimuli were achromatic gratings at the preferred spatial frequency and temporal frequency, at a contrast of 64% or greater. All stimuli were presented in a circular window confined to the classical receptive field (CRF). The measure of orientation selectivity that we used in this study was the bandwidth, which is the width of the smoothed response tuning using the width at 1/sqrt(2) (Schiller et al, 1976a; Ringach et al, 2002). Apart from the use of F0 responses for neurons with F1/F0 ratios < 1 and the F1 response for neurons with F1/F0 ratios > 1, details are the same as those given in a previous study (Ringach et al, 2002). Direction selectivity (DI – direction index) was determined from the orientation tuning as the response to the preferred orientation and drift-direction minus the response to the preferred orientation and opposite drift-direction divided by the sum of these two responses.

#### Spatial frequency tuning

Each neuron was presented with a range of spatial frequencies, usually in ½ octave steps from below 1 c/deg to around 10 c/deg. For some neurons the upper limit was extended if the neurons had responses to higher spatial frequencies. For almost all neurons that were orientation selective we measured spatial frequency tuning at the preferred orientation and drift-direction as well as at the non-preferred drift direction (Figs. 1-3 B, G, L, Q, see pairs of filled and unfilled circles). Each set of tuning responses was fit with a difference of offset difference of Gaussians (d-DOG-s; Hawken and Parker, 1987; Parker and Hawken, 1988) that provides a smooth fit to the data. We obtained the peak spatial frequency and the spatial frequency bandwidth (Tolhurst and Thompson, 1981; width/peak) from the fitted functions. For neurons where the response at low spatial frequency (0.1 c/deg) in the fitted function did not go below ½ the peak we called those cells low-pass (lp) in the figures but gave them bandwidth values of 6 in the cluster analysis (see Cluster Analysis Method).

**Figure 1:**
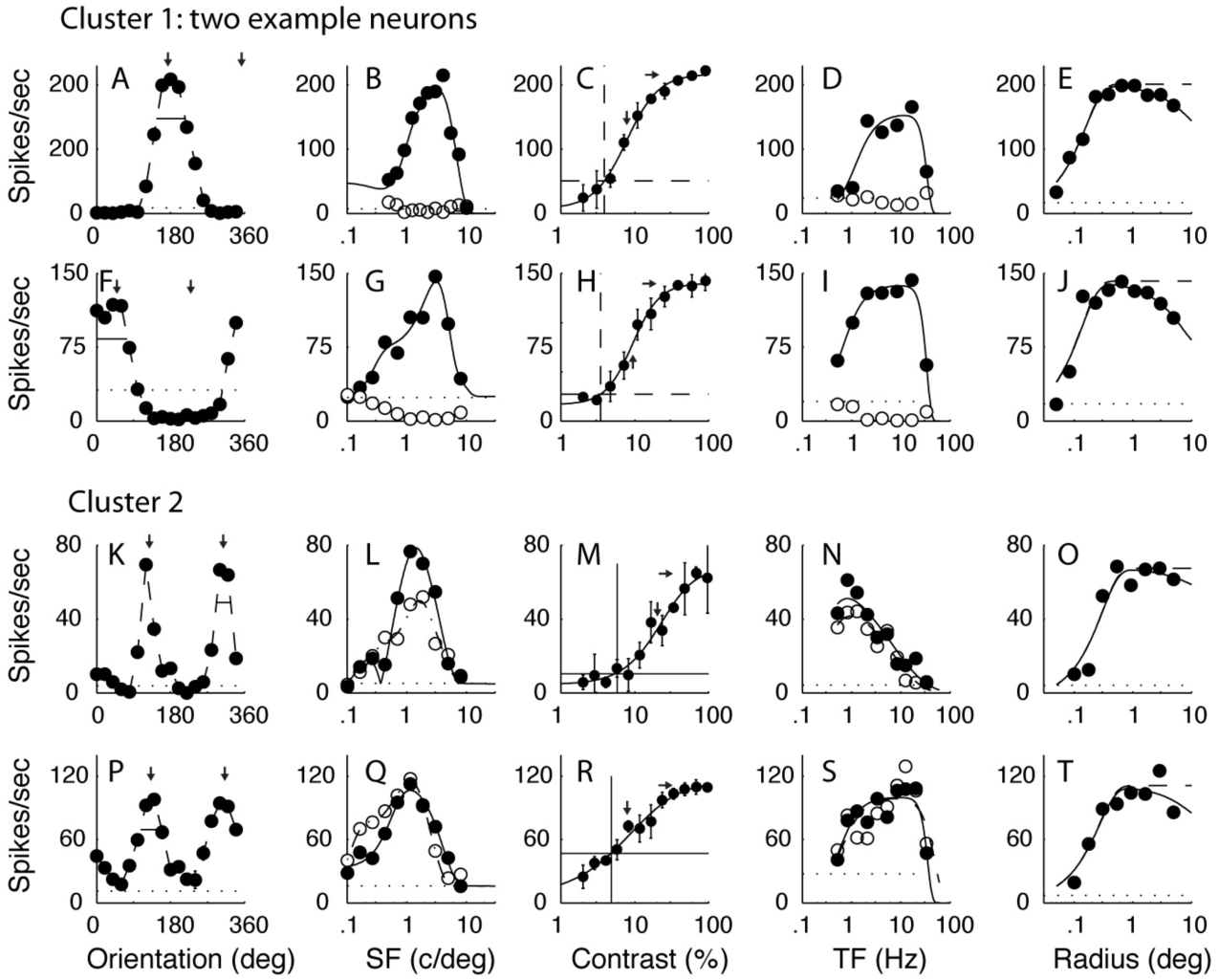
The tuning functions of example neurons from the two main clusters that had f1/f0 ratios < 1. A-E show the tuning from a single neuron in C1 (f1/f0 = 0.17). F-J show the tuning for a second neuron in C1 (f1/f0 = 0.07). The neurons in C1 were orientation and direction selective, had low contrast thresholds and were bandpass in temporal frequency. Also note the high response rate of these neurons which was a characteristic of this cluster. K-O show the tuning for a neuron in C2 (f1/f0 = 0.19). P-T show the tuning for a second neuron in C2 (f1 /f0 = 0.24). Neurons in this cluster were orientation selective but not direction selective, and were mainly low pass in temporal frequency (N) but some were bandpass (S). In A,F,K,P the response of each neuron was measured as a function of orientation to a drifting achromatic grating at the optimal spatial and temporal frequency and a high contrast through a window of the optimal size. The pair of small vertical arrows in each plot indicate the optimal orientation that was selected by fitting a von Mises function to the data, the small horizontal bars show the bandwidth – width at amplitude of √2 of the maximum. In B,G,L,Q the response of each neuron was measured as a function of spatial frequency at the preferred orientation and in both drift directions. The pairs of data in each plot denote the response at the preferred orientation for each drift direction (filled circles preferred drift direction, unfilled circles non-preferred drift direction) and the smooth curves fitted to the data are a spatial receptive field model that is a difference of offset difference-of-Gaussians (see Methods). In C,H,M,R the response as a function of contrast was measured for each neuron at the preferred orientation, drift direction, spatial and temporal frequency in a patch of grating of the optimal diameter. The smooth curve through each data set is a modified Naka-Rushton function (see Methods). The horizontal dotted line shows the response that is two standard deviations above the response to a blank grey screen (spontaneous response) and the vertical dotted line gives the value of contrast that was taken as the contrast threshold, the point where the horizontal line intersected the fitted Naka-Rushton function. In D,I,N,S the response as a function of temporal frequency is shown for a grating of the optimal orientation, spatial frequency and in both the preferred and opposite to preferred direction of drift direction (filled circles preferred drift direction, unfilled circles non-preferred drift direction). The two data sets were fitted independently with a difference-of-exponentials function and the best fitting functions are shown by the smooth curves. In E,J,O,T a grating of the optimal orientation, drift direction, spatial and temporal frequency and at high contrast (>= 64%) was presented in a circular window of different radii. The response as a function of radius of the window is shown and the data is fitted with a difference of Gaussians function (see Methods).

**Figure 2:**
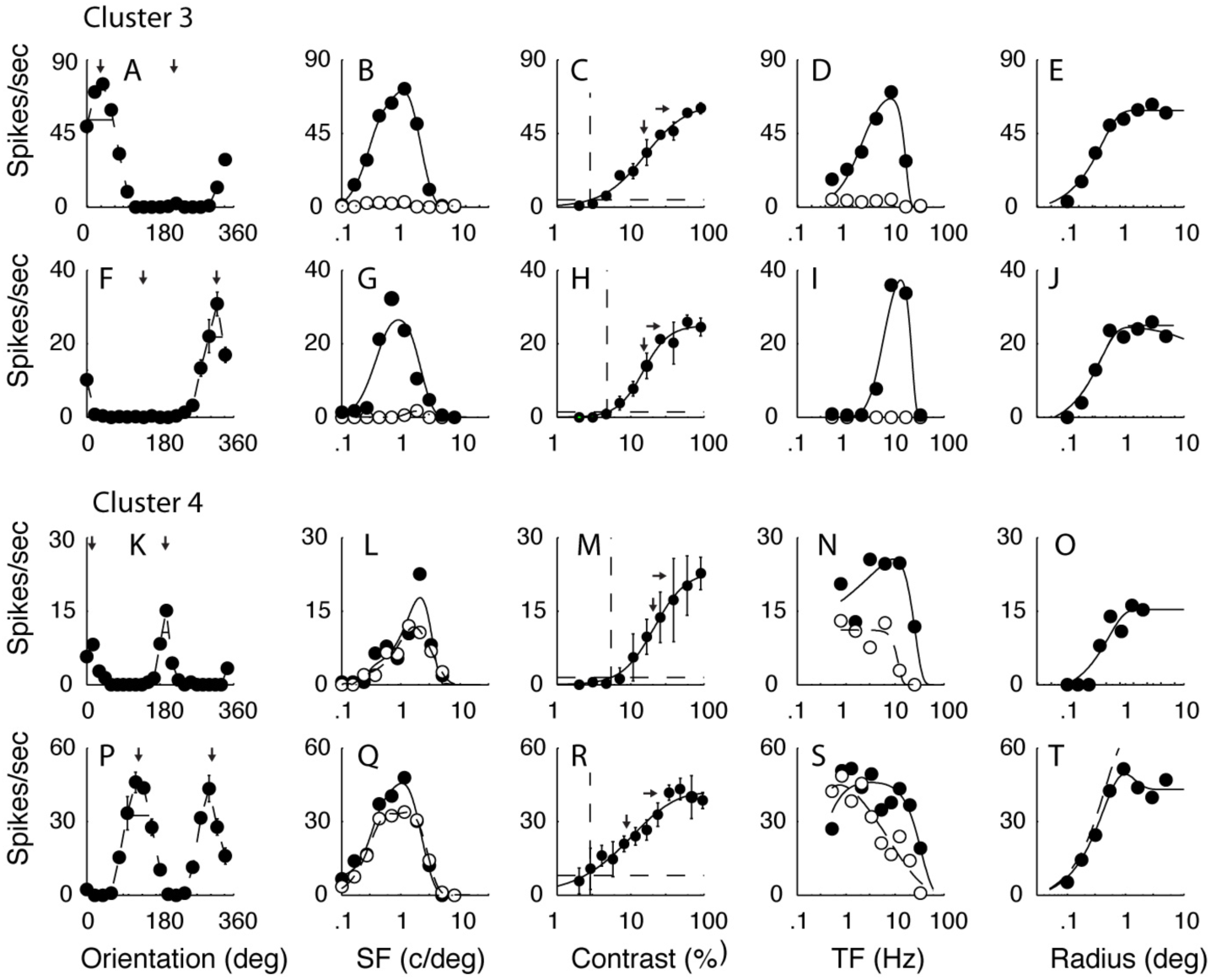
The tuning functions of example neurons from the two of the four main clusters that had f1/f0 ratios > 1. A-E and F-J are the tuning of two example neurons from C3 (f1/f0 1.5 and 1.1 respectively). Note that these neurons, characteristic of the cluster, were strongly direction selective and bandpass in temporal frequency. K-O and P-T are tuning functions of two example neurons from C4 (f1/f0 1.6 and 1.7 respectively), they were orientation but not direction selective with lowpass temporal frequency tuning. The other details are as for Fig. 1.

**Figure 3:**
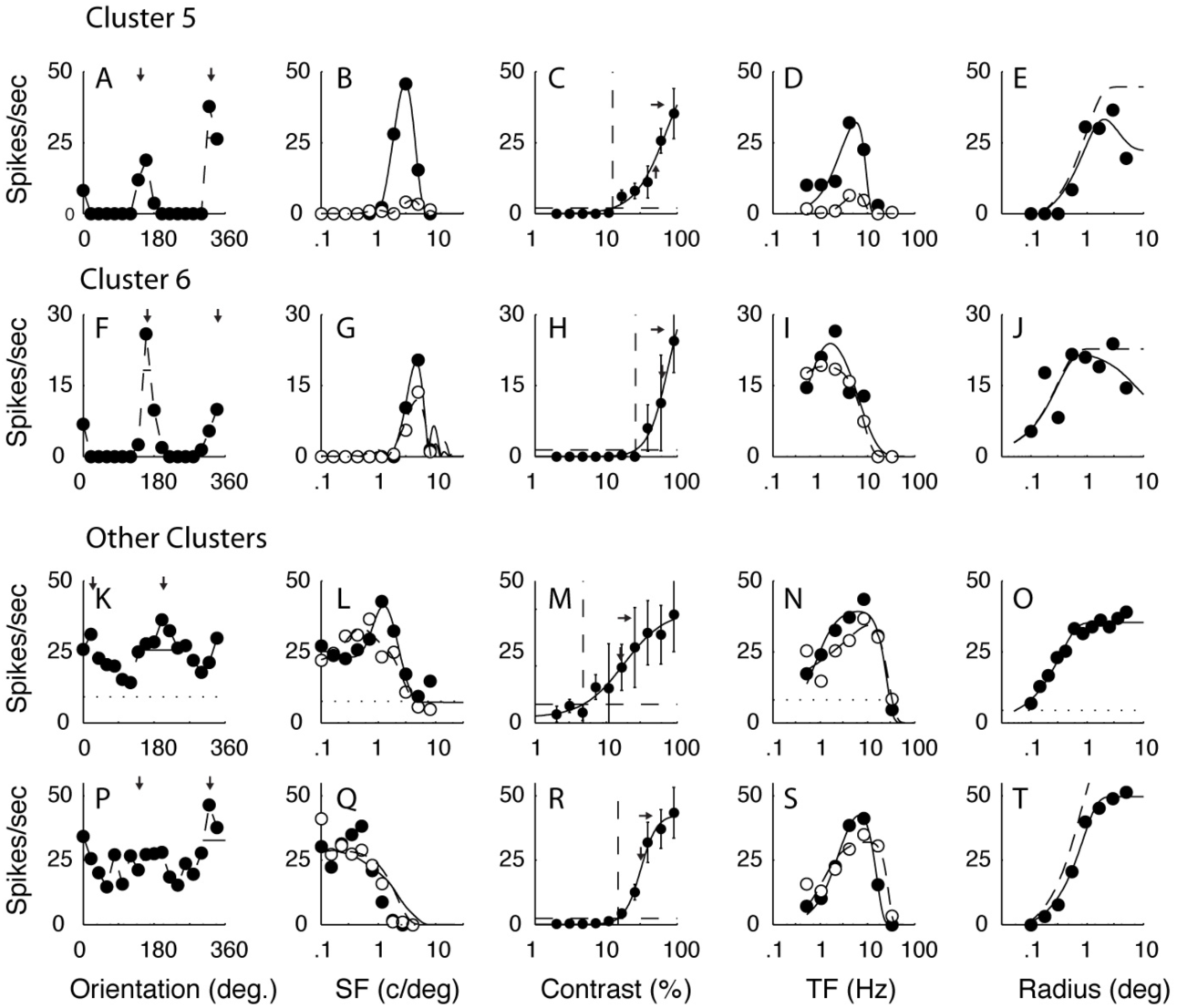
A-E, F-J show the tuning functions of example neurons from C5 and C6. The neurons in these clusters were orientation and spatial frequency selective, but one characteristic that separated them from the other two clusters was their insensitivity to contrast. They are separated from each other by their temporal frequency tuning, neurons in C5 were bandpass and those in C6 were lowpass. K-O and P-Q show neurons from the smaller clusters where most of the neurons were weakly selective for orientation and relatively lowpass in spatial frequency (f1 /f0 1.3 and 1.5 respectively). The other details are as for Fig. 1.

#### Contrast response

The response as a function of contrast was determined for contrasts ranging from 2% to 100% in half octave steps (Figs. 1-3 C, H, M, R). A blank condition (screen of mean grey luminance) of the same duration as the stimulus presentation was shown at the beginning of each sequence and interleaved between each contrast presentation. The contrasts were shown in ascending order to avoid adaptation and to minimize hysteresis (Bonds 1991). Each grating presentation lasted for a minimum of 4 temporal cycles, often more, and was at least one second in duration. Each dataset was fit with a modified Naka-Rushton function (Peirce, 2007) that captured the decrement in response (supersaturation) observed in the contrast response data of some V1 neurons. Two parameters from the fitted contrast response were obtained for each neuron: the contrast threshold, and the contrast (c50) at which the response reached 50% of the maximum rate. Note that the c50 was determined from the fitted curve, it was not the parameter in the modified Naka-Rushton – the so-called c50 parameter in the equation. The contrast threshold (cTh) was defined as the contrast where the fitted function just exceeded a threshold criterion, the measured spontaneous rate plus two standard deviations of the spontaneous rate. The threshold criterion is drawn as a horizontal dashed line in each example (Figs 1-3, C, H, M, R). The point where the criterion intersects the fitted function is the contrast threshold that was used in the current study. The vertical dashed line in each graph marks the threshold. The right-most short horizontal arrow in each plot is the maximum response of the fitted function. c50 is the point where response reached half the difference between the spontaneous rate and the maximum rate, and it is indicated by the short vertical arrow.

#### Temporal frequency tuning

Each neuron was presented with a range of temporal frequencies, usually in 1 octave steps from 0.5 Hz to 32 Hz. Measurements were made for drifting gratings at the optimal orientation and in the preferred and non-preferred direction shown by the pairs of symbols in Figs 1-3 ((Figs. 1-3 D, I, N, S; see pairs of filled and unfilled circles). Each set of responses was fit with a difference of exponentials function (Hawken et al, 1996), and from the fitted function to the preferred drift direction we obtained the temporal frequency at the maximum response amplitude, the peak TF. In addition we obtained the TF bandwidth from the preferred direction fitted function, defined as the width of the tuning function when the response was 50% of the maximum. Neurons where the response at low temporal frequency (0.1 Hz) in the fitted function did not go below ½ the peak were called low-pass (lp) in the figures but were given bandwidth values of 6 in the cluster analysis (see Cluster Analysis Methods).

#### Area summation

The response was measured as a function of stimulus area for a circular patch of grating of optimal spatial frequency, temporal frequency, orientation and direction of drift at a contrast of about 90% of the contrast that produced the maximum response (Fig. 1B). Initially the center of the CRF was estimated by moving an optimal drifting grating with a small aperture (usually 0.1 to 0.2 degrees in radius) until the experimenter judged that the position of maximum discharge rate had been reached. The patches of grating were centered on the x-y coordinates of the CRF center. A range of sizes from 0.05 or 0.1 degree up to 4 degrees radius was shown in half octave steps. Stimuli were shown in pseudorandom order for each repeat of the sequence. The number of repeats was at least three. A blank stimulus of a mean grey screen of the same duration as the stimulus was shown between each repeat. The responses were fit with a difference of Gaussians model (Sceniak et al, 1999; 2001) and the best fitting functions are shown as the smooth blue curves in Figs. 1 – 3 (E, J, O, T). A measure of area suppression (SI – suppression index) was obtained from the area tuning as described in Sceniak et al (2001).

#### Peak and spontaneous firing rate

The peak firing rate was obtained from each tuning curve. For example in the orientation domain the peak rate was the orientation and drift rate that evoked the maximum response. For neurons with f1/f0 < 1 (complex cells) the rate was calculated as the average rate across the full duration of the stimulus presentation (f0) while for neurons with f1/f0 >= 1 the rate was the amplitude of the first harmonic response (f1). The spontaneous rate for all neurons was the mean rate during the blank (mean grey screen) intervals that were the same duration as the stimulus intervals and that were interleaved with the stimulus presentation.

### Cluster Analysis Method

Seven response measures and tuning parameters obtained from orientation tuning, spatial and temporal frequency tuning, and contrast-response measurements were used as the input parameters to the cluster analysis. The parameters used for clustering were f1/f0 ratio, orientation bandwidth, direction index, spatial frequency bandwidth, temporal frequency bandwidth, contrast threshold and c50 contrast. Each value was converted to a z-score for the corresponding distribution. For the three bandwidth measures, if the neuron’s tuning was unoriented (in that the response did not drop below 50% of the maximum response) or low pass in spatial or temporal frequency we used values of 120 degrees (orientation bandwidth), and 6 octaves (spatial frequency and temporal frequency bandwidth) as the values for calculating the z-scores. These values are likely to be lower than the real bandwidths of effectively lowpass neurons and thus are conservative values. Nonetheless they allowed us to include the lowpass or unoriented tuning functions in the clustering determination. These values are shown as lowpass (lp) in the tuning distributions.

The z-scores were input to the Matlab functions pdist and linkage. In linkage, the standard Euclidean distance metric and the unweighted average distance were used in the linkage-clustering algorithm. The output of linkage was used to obtain the hierarchical clusters. With a large set of tuning and response measures, finding the unique optimal number of clusters is not guaranteed. The silhouette function (Matlab) was applied to the output of the linkage routine to estimate the number of clusters. The silhouette value is determined for each point to measure the distance of that point from its assigned cluster compared to points in other clusters. The value ranges from – 1 to 1 where a value close to 1 means the assignment is appropriate. The accumulated sum of silhouette values increases if adding additional clusters provides improved clustering until adding more doesn’t make much difference i.e. most of the additional points are ∼0 and therefore the accumulated value plateaus - the first plateau can be used as an approximate guide as to the maximum number of clusters to specify in the functions pdist and linkage. In the layer 6 dataset the plateau was reached with 22 clusters. Six of the clusters contained the majority of the population (N = 85/116, 73%), and appeared to divide the population into distinct clusters.

Statistical comparisons between parameters in clusters were made using a multiple comparison of the output of a one-way ANOVA using the Bonferroni method to provide the adjustment to compensate for multiple comparisons. The p-values in Results for all comparisons are shown in Tables 1-9. The tests do not test for the significance of clusters but evaluate whether the parameters were significantly different between clusters.

**Table 1:** Orientation Bandwidth: The values in the first column under the cluster number show the mean ± 1SD for each cluster. The p values (below the diagonal) were all determined using a multiple comparison of the output of a one-way ANOVA using the Bonferroni method to provide the adjustment to compensate for multiple comparisons. Values above the diagonal are 1 – log(p) indicating the full range of significance values. All values shown as bold reached significance level < 0.05.

|  | **C1** | **C2** | **C3** | **C4** | **C5** | **C6** |
| --- | --- | --- | --- | --- | --- | --- |
| **C1**<br>29.1 ± 11.4 |  | 1 | 7.0 | 4.9 | 5.1 | 3.5 |
| **C2**<br>26.9 ± 12.4 | p = 1 |  | 5.6 | 3.5 | 3.9 | 1.3 |
| **C3**<br>15.9 ± 5.3 | **p < 0.003** | **p < 0.01** |  | 1 | 1 | 1 |
| **C4**<br>17.5 ± 9.1 | **p < 0.03** | p = 0.08 | p = 1 |  | 1 | 1 |
| **C5**<br>15.4 ± 4.5 | **p < 0.02** | p = 0.06 | p = 1 | p = 1 |  | 1 |
| **C6**<br>18.9 ± 10.5 | p = 0.27 | p = 0.76 | p = 1 | p = 1 | p = 1 |  |

## Results

We applied cluster analysis to a population of 116 neurons that we assigned to layer 6. The methods of layer assignment were described previously (Hawken et al, 1988; Ringach et al, 2002). As described in Methods, in the cluster analysis we used seven tuning parameters: f1/f0 ratio, orientation bandwidth, directional index, spatial frequency bandwidth, temporal frequency bandwidth, contrast threshold, C50 contrast. The clustering returned four clusters (C1 – C4) that each had ten percent or more of the neurons and together accounted for 59% of the population. Two smaller clusters (C5 and C6) had 8% and 7% of the neurons. It is particularly informative that there were other visual response properties, which were not used as data in the clustering analysis, but were distributed according to the clustering we obtained. Those data will be presented after first examining the tuning curves of clustered neurons on the dimensions used for clustering.

### Single cell examples

#### Tuning examples in clusters

First, we show the distinctive tuning within each of the clusters by presenting examples of the tuning curves of neurons in the six major clusters in the dimensions of stimuli used in the clustering analysis and compare tuning curves between neurons in the major clusters. Also, we present a few examples of tuning curves of neurons from the smaller clusters.

The first main cluster (C1: n = 15/116, 13%) was comprised of highly responsive and highly sensitive, direction-selective complex cells (two examples in Figure 1A-E, and F-J). One distinguishing parameter in the cluster analysis was the f1/f0 ratio. The f1/f0 ratios of the two example neurons from the first main cluster were 0.17 and 0.07, both in the lower quartile of ratios of complex cells. Another major distinguishing feature of the tuning of the first cluster was that all neurons were strongly direction-selective (Fig. 1A, F). Neurons in the first cluster often showed strong suppression (below the spontaneous baseline shown by the horizontal dotted line) in the non-preferred direction for a grating of the optimal orientation (e.g. Fig. 1F). Many of the neurons in C1 also had relatively low contrast thresholds and saturating contrast response functions (Fig. 1C, H). In the temporal frequency domain, most of the neurons in C1 were strongly band-pass in temporal frequency tuning as exemplified in the two example neurons (Fig. 1D, I). In the experiment where the radius of the stimulus window was changed, some neurons in C1 showed a modest attenuation of their response at the largest window sizes (Fig. 1E,J), although no parameter from this experiment was used in the cluster analysis.

The neurons in the second major cluster (C2: n = 20/116, 17%) also had low f1/f0 ratios (0.19, 0.24 for the two examples shown in the third and fourth rows of Fig 1). The neurons in C2 were orientation-selective but not direction-selective (Fig. 1K, P), unlike those in C1. There were some neurons in C2 that were low-pass in temporal frequency (Fig. 1N) and others that were band-pass (Fig. 1S). Neurons in C2 showed very little size tuning (Fig. 1O, T).

The neurons in the other four major clusters (C3 – C6) had f1/f0 ratios greater than 1, and therefore were classified as simple cells (Skottun et al, 1991). Ratios for the example neurons in C3 (Fig. 2A-J), C4 (Fig. 2K-T), C5 (Fig. 3A-E) and C6 (Fig. 3F-J) were 1.5, 1.1, 1.6, 1.5, 1.4 and 1.8 respectively. The example simple cells in C3 (n = 18/116, 16%) showed orientation selectivity and strong direction-selectivity (Fig. 2A, F) that were characteristics of this cluster. The strong directional-selectivity was observed across all spatial (Fig. 2B, G) and temporal (Fig. 2D, I) frequencies tested for the two example neurons. Another distinguishing tuning characteristic of the third cluster is shown by the band-pass temporal frequency tuning of the two example neurons (Fig. 2D, I); band-pass temporal frequency tuning was also a common feature of this cluster.

The neurons in C4 (n = 15/116, 13%) were orientation-selective but not direction-selective (Fig. 2K, P). The lack of direction-selectivity in C4 was evident in the spatial (Fig. 2L,Q) and temporal (Fig. 2N,S) frequency tuning of the example neurons. The tuning shape for both spatial (Fig. 2L, Q) and temporal (Fig. 2N, S) frequency along with the response amplitude was similar in both directions of movement at the preferred orientation. The temporal frequency tuning of the two example neurons from cluster 4 were low-pass. This was a common but not universal feature of C4. Most of the neurons in clusters 3 and 4 showed little attenuation in their response at large stimulus sizes as shown for the four example neurons (Fig. 2E, J, O, T).

One distinguishing feature of two additional clusters (C5, n = 9 and C6, n = 8) with f1/f0 ratios > 1 was their low sensitivity to contrast as can be seen in the non-saturating contrast response functions from the two example neurons (Fig. 3C, H). Neurons in C5 and C6 were orientation-selective, and some were moderately direction-selective (Fig. 3A). However, they were distinguished by their temporal frequency tuning: neurons in C5 were bandpass in temporal frequency (Fig. 3D) while those in C6 were lowpass in temporal frequency (Fig. 3I).

The remaining clusters had few members (5 or fewer) and therefore it was not possible to verify their uniqueness with the relatively small population sample. Two features separated most members of these relatively small clusters from the six larger clusters. One was broad orientation tuning. Often neurons had a preferred orientation but also responded across all orientations with a response at the orthogonal-to-preferred orientation (Fig. 3K, P). The second distinguishing feature was that the broad orientation tuning was often accompanied by weak-bandpass or lowpass spatial frequency tuning (Fig. 3L, Q).

### Population analyses

We studied how the seven parameters that were used in the clustering were distributed between the different major clusters. Following this we also studied how surround suppression, peak response rate and spontaneous rate varied between the clusters.

#### F1/F0 distribution – Simple and Complex Cells

The f1/f0 ratio was used as a clustering parameter. f1/f0 is the ratio of the first-harmonic (f1) modulation amplitude of the response to the elevation of response (f0). This is a straightforward method of distinguishing simple (f1/f0 ratio > 1) from complex cells (f1/f0 ratio < 1) (Campbell & Fiorentini 1973; Movshon et al, 1978; DeValois et al, 1982a,b; Hawken & Parker, 1987; Skottun et al, 1991). All the major clusters that were identified had neurons with either f1/f0 < 1 (clusters 1, 2) or f1/f0 > 1 (clusters 3, 4, 5, 6). The distribution of f1/f0 for the population of layer 6 neurons was bimodal with a very sharp null close to 1 (Fig. 4A).

**Figure 4:**
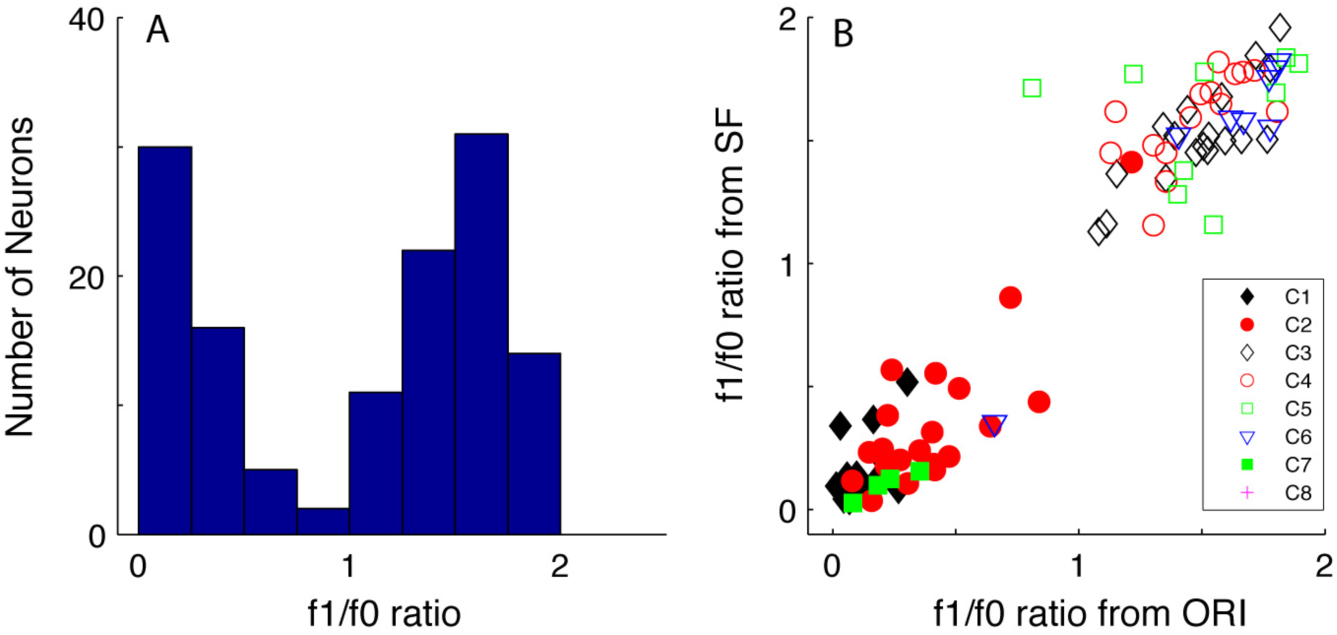
A.The distribution of f1/f0 ratio for all the neurons in the sample from layer 6 (n = 116).The f1/f0 ratio was used as one of the response measures in the clustering procedure. B: shows the f1/f0 ratio measured in the orientation tuning experiment (x-axis) plotted against the f1/f0 ratio of the same cell measured from the spatial frequency tuning experiment (y-axis). Each symbol denotes a cluster. There were six main clusters that are shown as filled black diamonds (C1), filled red circles (C2), unfilled black squares (C3), unfilled red circles (C4), unfilled green squares (C5) and unfilled blue triangles (C6).Three smaller clusters with f1/f0 ratios > 1 are shown with by unfilled magenta triangles and cyan and magenta crosses and a small cluster with f1/f0 < 1 is shown by the filled blue triangles. There were four further clusters with f1/f0 < 1 and with fewer than 4 neurons that were combined and are shown as filled green squares. The clusters were clearly separated according to whether they are complex (f1/f0 < 1) or simple (f1/f0 >1).

The f1/f0 ratio was relatively consistent when determined for the same neuron in different experiments (Fig. 4B). Neurons with f1/f0 > 1 (classified as simple) in the orientation tuning experiment (Fig. 4B x-axis) remained so in the subsequent spatial frequency tuning experiment (Fig. 4B y-axis). Only one of the 85/116 neurons in the major clusters crossed the S/Cx division between experiments – it was classified as a complex cell but segregated with a simple cell cluster. It was a cell with f1/f0 = 0.8 that was assigned to cluster 5 where all the other members had f1/f0 > 1. In the current data set, f1/f0 was obtained from the orientation tuning experiment. When we checked f1/f0 for this cell in other experiments, f1/f0 was > 1 (1.71 for SF, 1.61 for RVC, 1.65 for TF and 1.63 for area) so clearly it was on the border. Considering all experiments together, we classified it as a simple cell.

#### Direction Index and Orientation Bandwidth distributions

Clusters of cells in layer 6 were distinguished by their direction selectivity and also orientation selectivity estimated from orientation tuning bandwidth. Direction selectivity was determined from the direction index (dI see Methods). A value of dI near zero means no preference for direction and dI near 1 means almost completely direction selective. Neurons that are suppressed in the non-preferred direction can have a dI > 1.

The distribution of direction index, dI, in layer 6 was bimodal (Fig. 5B); 39% (41/104) of orientation selective neurons had a dI >= 0.8. There was a population of very direction-selective neurons in layer 6. Neurons in the orientation- and direction-selective complex cell cluster C1 (Fig. 5C, filled black diamonds) had an average dI of 1.05 ± 0.3 (mean ± 1sd) which is significantly different (p < 0.0001) from the dI of neurons in the non-direction selective complex cell cluster C2 (Fig. 5C, filled red circles) in which mean dI = 0.16 ± 0.14. Although C1 and C2 differed in dI, they did not have different orientation bandwidth distributions (29.1 ± 11.4 deg. Vs 26.9 ± 12.4 deg. for C1 and C2 respectively, p = 1, n.s.). The dI distributions of the simple cell orientation and direction-selective cluster, C3 (Fig. 5D, black diamonds), and the non-direction-selective clusters, C4 (Fig. 5D red circles), C5 (Fig. 5F green squares) and C6 (Fig. 5F blue diamonds) were significantly different (p < 0.0001, Table 2). All four simple cell clusters had relatively narrow oBW’s (Fig. 5D, F) that were not significantly different between clusters (Table 1).

**Table 2:** Direction Index: The values in the first column under the cluster number show the mean ± 1SD for each cluster. The p values were all determined as in Table 1.

|  | **C1** | **C2** | **C3** | **C4** | **C5** | **C6** |
| --- | --- | --- | --- | --- | --- | --- |
| **C1**<br>1.05 ± 0.30 |  | 39.9 | 1 | 26.8 | 19.1 | 23.1 |
| **C2**<br>0.16 ± 0.14 | **p < 0.0001** |  | 33.7 | 2.2 | 2.8 | 1 |
| **C3**<br>0.90 ± 0.13 | p = 0.99 | **p < 0.0001** |  | 20.1 | 13.2 | 17.4 |
| **C4**<br>0.34 ± 0.31 | **p < 0.0001** | p = 0.31 | **p < 0.0001** |  | 1 | 1 |
| **C5**<br>0.40 ± 0.24 | **p < 0.0001** | p = 0.17 | **p < 0.0001** | p = 1 |  | 1 |
| **C6**<br>0.28 ± 0.20 | **p < 0.0001** | p = 1 | **p < 0.0001** | p = 1 | p = 1 |  |

**Figure 5:**
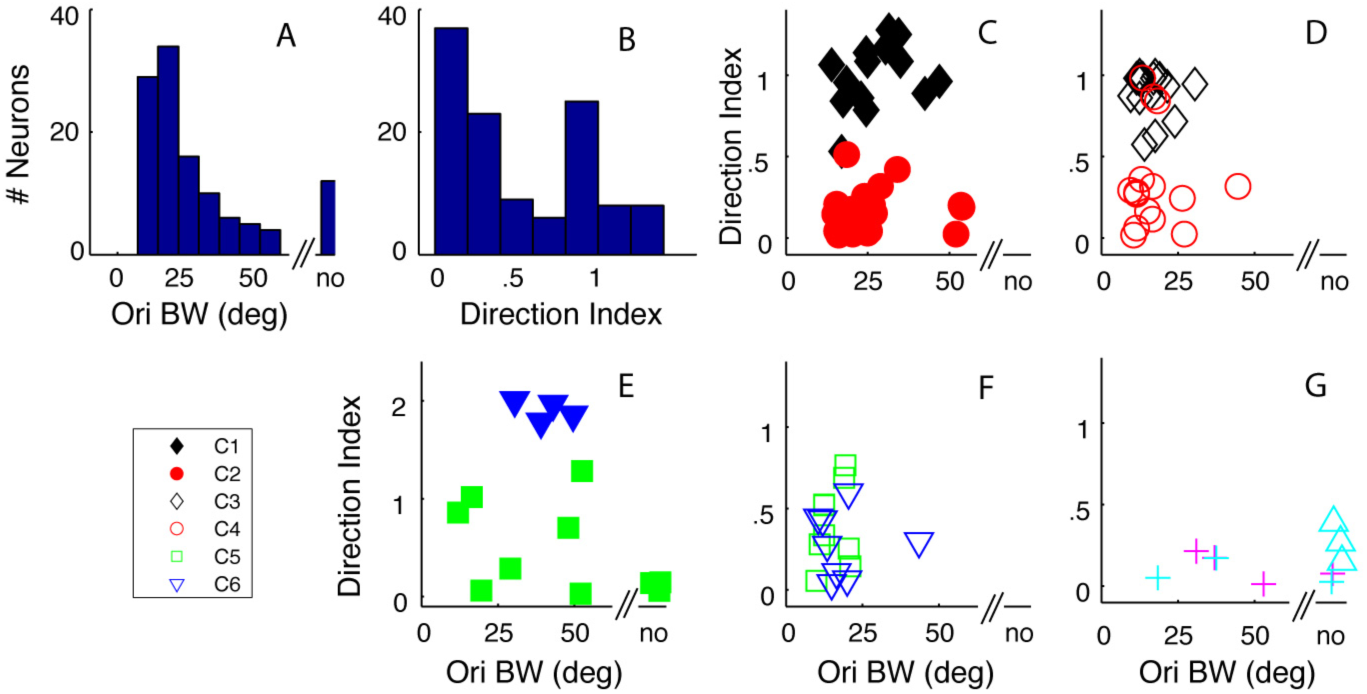
Orientation bandwidth (obw) and direction index (dl) were two tuning parameters used in the clustering procedure. A: shows the distribution of obw for the whole layer 6 sample, no - non-oriented. B: The distribution of dl for the whole sample. C-G: shows the relationship between obw (x-axis) and dl (y-axis) for the same clusters shown in Fig. 4. C: shows the obw/dl relationship for C1 (filled black diamonds) and C2 (filled red circles). There is a clear separation by dl but not obw in the two clusters. D: shows the relationship for C3 (unfilled black diamonds) and C4 (unfilled red circles). There is a clear separation based on dl but not obw in the two clusters. E: shows the obw/dl relationship between the remaining clusters where the clusters also had f1/f0 < 1..The neurons in the small cluster denoted by the filled blue triangles are very direction selective and strongly suppressed in the non-preferred direction. The filled green squares are the combination of three small clusters. F: shows the obw/dl relationship for C5 (green squares) and C6 (blue triangles), both with neurons that have f1/f0 > 1. G. three small clusters (unfilled cyan triangles, magenta crosses and cyan crosses where all the clusters also had fl/f0 < 1.

Most neurons we studied in layer 6 had a measurable orientation bandwidth although a small number (n = 12/116, 10%) were non-oriented (Fig. 5A). The distribution of orientation bandwidth for all orientation-selective neurons in layer 6 had a median value of 20.5 deg. (Fig. 5A). Although when comparing within the same f1/f0 clusters there were no significant differences in orientation bandwidth, there were significant differences between C1 and C3 – C5 and between C2 and C3. (see Table 1 for p values).

There were a number of smaller clusters (Fig. 5E, G) for which we did not test differences because of the small number of neurons in each cluster. However qualitatively, one cluster with f1/f0 < 1 had dI’s of around 2 (Fig. 5E blue filled triangles), these were distinct from all the other neurons as they had relatively high maintained rates and showed a large suppression in the opposite to preferred direction. There were other neurons with f1/f0 < 1 that formed small clusters with a range of dI and orientation bandwidths that are shown in Fig. 5E (green squares). Other neurons with f1/f0 > 1 that were in the minor clusters were relatively untuned for orientation as in the example neurons in Fig 3K,P and accounted for some neurons in the smaller clusters (Fig. 5G).

#### Spatial Frequency Tuning

One of the characteristic features of many cortical receptive fields is that they have multiple subunits (Hubel and Wiesel, 1962; 1968) that can result in relatively narrow spatial frequency tuning of neurons (Movshon et al, 1978; DeValois et al, 1982; Hawken and Parker, 1987). We measured the spatial frequency tuning at the preferred orientation, drift direction and temporal frequency for high contrast gratings (>= 64% contrast) and determined the bandwidth from the d-DOG-s function fitted to the response data (see Methods). All but two of the neurons in C1 were band-pass in spatial frequency (Fig. 6A, black diamonds); the population had mean bandwidth of 2.4 ± 1.2 octaves (mean ± 1SD) and was not significantly different (Table 3) from the bandwidth distribution of neurons in C2 (2.9 ± 1.3 octaves, mean ± 1SD) where two neurons were low-pass (Fig. 6A, red filled circles). All the neurons in the other four major clusters were band-pass in spatial frequency (Fig. 6B black unfilled diamonds – C3; red unfilled circles – C4; Fig. 6D green unfilled squares – C5 and blue triangles – C6). None of their spatial frequency bandwidth distributions were significantly different (Table 3). The spatial frequency bandwidth distribution of C6 was significantly narrower than the distribution from both C1 and C2, while the distributions from C3, C4 and C5 were narrower than C2 (Table 3). Some of the neurons in the smaller clusters were low-pass (Fig. 6C, E) while others were bandpass (Fig. 6C blue diamonds, Fig. 6E magenta crosses).

**Table 3:** Spatial Frequency Bandwidth: The values in the first column under the cluster number show the mean ± 1SD for each cluster. The p values were all determined as in Table 1.

|  | **C1** | **C2** | **C3** | **C4** | **C5** | **C6** |
| --- | --- | --- | --- | --- | --- | --- |
| **C1**<br>2.36 ± 1.19 |  | 1 | 1 | 1 | 3.0 | 4.2 |
| **C2**<br>2.93 ± 1.29 | p = 1 |  | 6.3 | 5.7 | 8.5 | 9.7 |
| **C3**<br>1.80 ± 0.64 | p = 1 | p < 0.005 |  | 1 | 1 | 1 |
| **C4**<br>1.79 ± 0.78 | p = 1 | p < 0.01 | p = 1 |  | 1 | 1 |
| **C5**<br>1.30 ± 0.34 | p = 0.13 | p < 0.001 | p = 1 | p = 1 |  | 1 |
| **C6**<br>1.10 ± 0.32 | p < 0.05 | p < 0.0005 | p = 1 | p = 1 | p = 1 |  |

**Figure 6:**
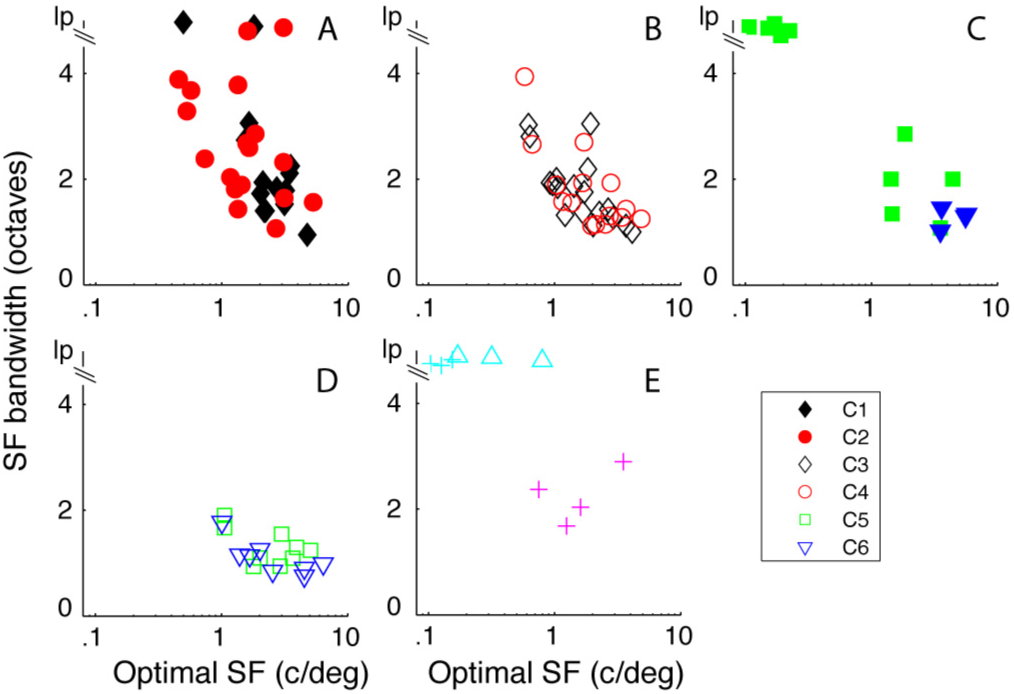
The relationship between optimal spatial frequency (osf, x-axis) and spatial frequency bandwidth (sfbw, y-axis) for the main clusters. Cluster designation as in Fig. 5. Ip - lowpass. A: C1 and C2, there in a inverse relationship between osf and sfbw for both groups. B: C3 and C4 show overlapping relationships between osf and sfbw. C: The small very direction selective cluster (filled blue triangles) have very similar osf’s and sfbws whereas the combined clusters have a number of neurons that are lowpass in spatial frequency tuning. D: C5 and C6 show narrow sfbws across a range of osfs. E: Neurons in two clusters are lowpass in spatial frequency, the other is bandpass (magenta crosses).

#### Temporal Frequency Tuning

Previous studies have shown that there is a range of temporal frequency tuning in V1. However, there has not been a systematic analysis at the level of single cortical layers. In the examples from the different layer 6 clusters, some neurons were strongly band-pass in TF (Fig. 1D, I; Fig. 2 D, I; Fig. 3D) while other neurons were low-pass in TF (Fig. 1N; Fig. 2N, S; Fig. 3I). We used temporal frequency bandwidth to quantify the width of the tuning and the degree of attenuation at low temporal frequencies. The high and low frequency half-height points were determined for each neuron from the fit of a difference-of-exponentials to the data (see Methods). If a fit did not attenuate to half amplitude by a TF of 0.1, we assigned the low TF half-height point = 0.1, then used this point and the high frequency at half amplitude to determine bandwidth. A number of neurons with no or weak and shallow low-frequency attenuation would have a slightly narrower bandwidth than their true bandwidth using this assignment, but we used it to allow us to make quantitative comparisons of the different clusters. Neurons with bandwidths > 6 are shown as lowpass (lp) in Figure 7.

**Figure 7:**
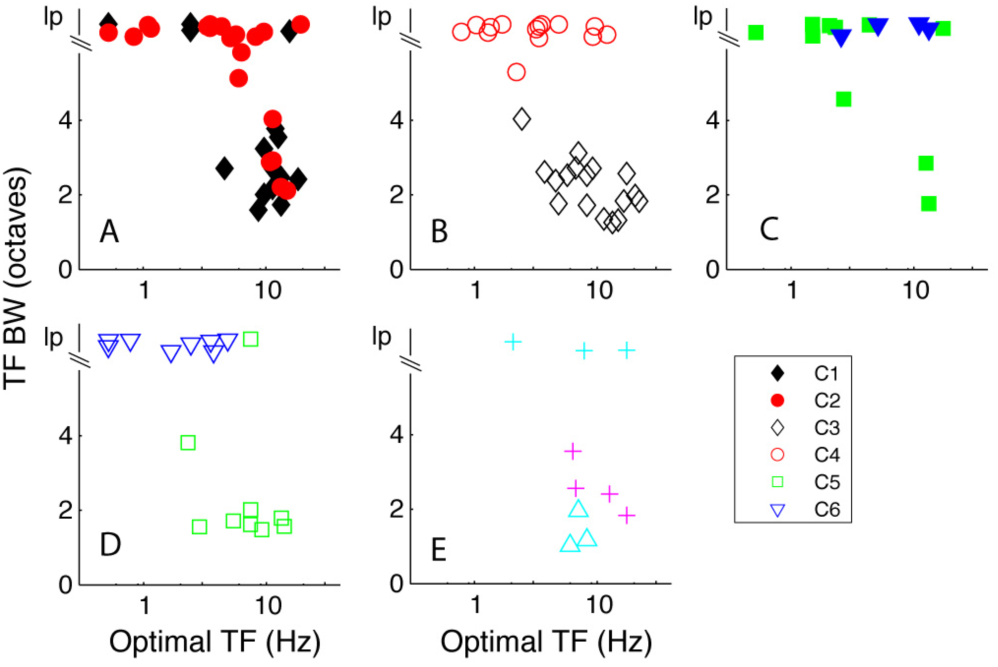
The relationship between optimal temporal frequency (otf, x-axis) and temporal frequency bandwidth (tfbw, y-axis) for the main clusters. Cluster designation as in Fig. 5. Ip - lowpass. A: C1 and C2, the majority of neurons in Cl are bandpass in temporal frequency and have otfs around 10 Hz. Many of the neurons in cluster 2 are Ip in temporal frequency. B: C3 and C4 show distinct separation based on tfbw. C:The majority of the neurons in the remaining clusters with f1/f0 ratio < 1 are lowpass in temporal frequency. D: C5 and C6 are separated by their tuning for TF (except 1 neuron in C5 that is Ip). E:Three small clusters are distinguished by their temporal frequency tuning.

Neurons in the main direction-selective complex cell cluster – C1 – were predominantly band-pass in TF (Fig. 7A, black filled diamonds) whereas the majority of the non-direction selective complex cell cluster – C2 – were mostly low-pass in their TF tuning (Fig. 7A, red filled circles). The mean bandwidths of the two main complex cell clusters C1 and C2 were significantly different (3.6 ± 1.9 vs 5.4 ± 1.6, p < 0.005, Table 4). There also was a clear separation in the TF bandwidths of the neurons in the two largest simple cell clusters C3 and C4 (Fig. 7B, black unfilled diamonds and red unfilled circles). Neurons in the direction-selective cluster – C3 – were all bandpass in TF while neurons in the non-direction selective cluster – C4 – were nearly all low-pass; the difference between these two clusters was significant (p < 0.0001, Table 4), as was the difference between C4 and C5 (p < 0.0001). However, temporal frequency bandwidth distributions of C4 and C6 were predominantly lowpass (Fig. 7B red circles and Fig. 7D blue triangles) and not significantly different (see Table 4). C5 and C6 had significantly different TF bandwidth distributions (Fig. 7D, Table 4).

**Table 4:**
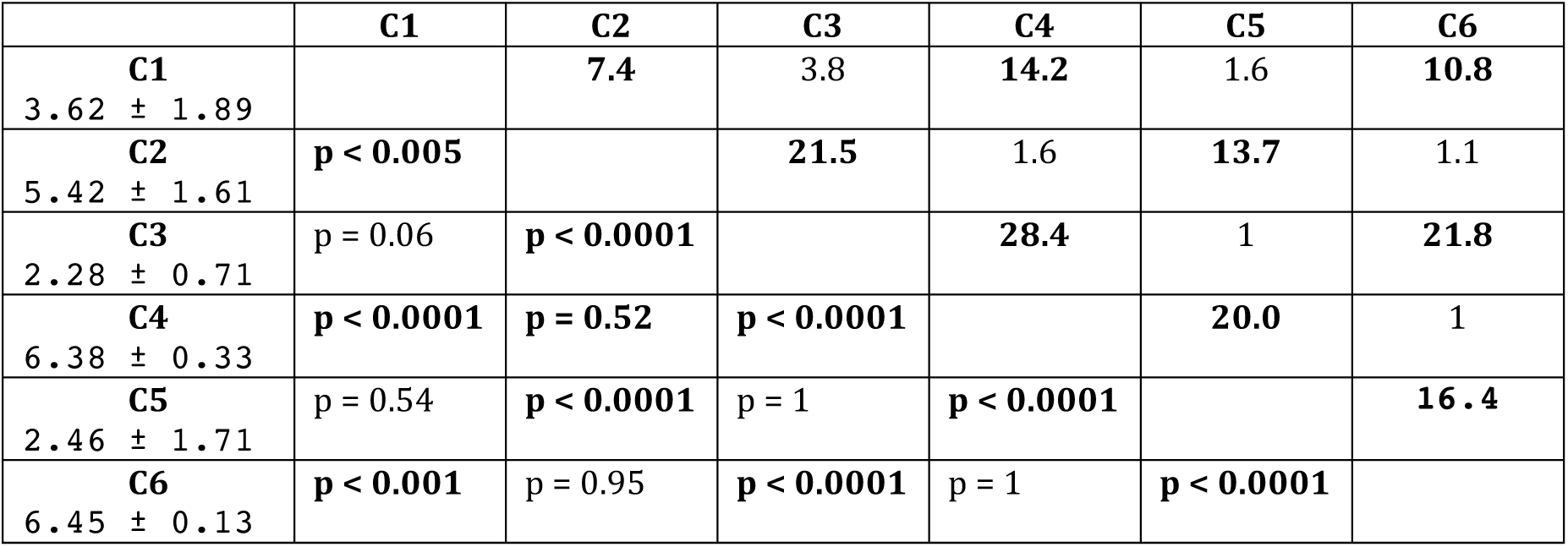
Temporal Frequency Bandwidth: The values in the first column under the cluster number show the mean ± 1SD for each cluster. The p values were all determined as in Table 1.

#### Contrast response distributions – threshold and c50

Each neuron’s response as a function of contrast for gratings of optimal orientation, direction, spatial and temporal frequency was fit with a modified Naka-Rushton function (Peirce, 2007; see Methods). Two measures of the response were obtained from the fitted functions and used as parameters in the clustering: 1) the contrast at which the response reached half its maximum (c_50_) and 2) the contrast threshold (c_Th_), defined as the contrast where the response exceeded twice the standard deviation of the baseline response. The contrast threshold parameter distributions from clusters C1-C4 (Fig. 8A, B) were not significantly different from each other (Table 5) but were all significantly lower than the parameter distributions from C5 and C6 (Fig. 8D, Table 5).

**Table 5:**
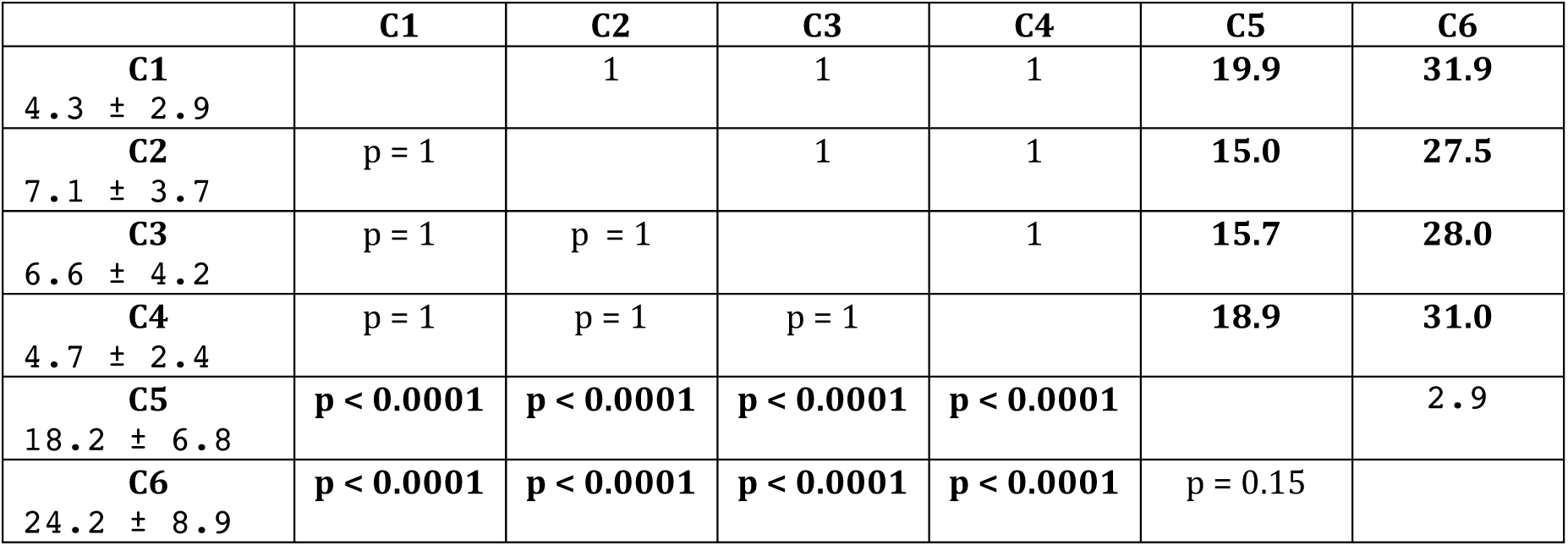
Contrast Threshold: The values in the first column under the cluster number show the mean ± 1SD for each cluster. The p values were all determined as in Table 1.

|  | **C1** | **C2** | **C3** | **C4** | **C5** | **C6** |
| --- | --- | --- | --- | --- | --- | --- |
| **C1**<br>4.3 ± 2.9 |  | 1 | 1 | 1 | **19.9** | **31.9** |
| **C2**<br>7.1 ± 3.7 | p = 1 |  | 1 | 1 | **15.0** | **27.5** |
| **C3**<br>6.6 ± 4.2 | p = 1 | p = 1 |  | 1 | **15.7** | **28.0** |
| **C4**<br>4.7 ± 2.4 | p = 1 | p = 1 | p = 1 |  | **18.9** | **31.0** |
| **C5**<br>18.2 ± 6.8 | **p < 0.0001** | **p < 0.0001** | **p < 0.0001** | **p < 0.0001** |  | 2.9 |
| **C6**<br>24.2 ± 8.9 | **p < 0.0001** | **p < 0.0001** | **p < 0.0001** | **p < 0.0001** | p = 0.15 |  |

**Figure 8:**
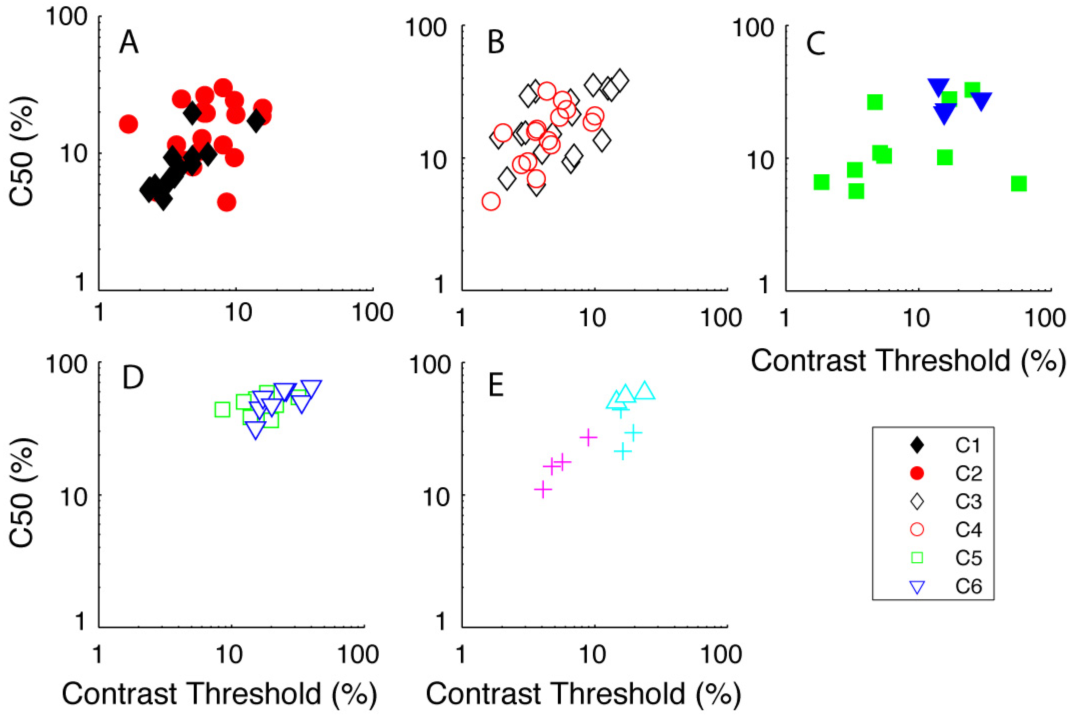
Two measures from the response as a function of contrast were used as tuning parameters in the clustering procedure: contrast threshold (cTh) and the contrast where the response reached half the maximum response (c50). Cluster designation as in Fig. 5. A: C1 has neurons with relatively low cTh and c50 whereas neurons in C2 are intermediate in their contrast sensitivity. B: C3 and C4 have overlapping cThs and c50s but are somewhat less sensitive than C1. C: Neurons in the direction selective cluster (blue triangles) are relatively insensitive to contrast. D: Neurons in C5 and C6 have high cThs and c50, making the clusters stand out as being contrast insensitive. E: One clusters is intermediate in contrast sensitivity (magenta crosses), another insensitive (cyan triangles).

A similar pattern was found for the c_50_. The neurons in C5 and C6 (Fig. 8D, green squares and blue diamonds) were relatively insensitive to contrast as seen in the example neurons (Fig. 3C, H). The mean values of the of c_50_’s parameter distributions for C5 and C6 (48.9 ± 8.2% and 52.2 ± 11.1% respectively) were not significantly different from each other (Table 6) but they were significantly higher than those of all the other main clusters (p < 0.0001, Table 6). There was relatively little overlap in the thresholds and c_50_ of C5 and C6 with the other four principal clusters. Among the other four clusters only the c_50_ parameter distribution for C3 was significantly higher than for C1 (Table 6). The smaller simple cell clusters had restricted ranges of c_50_ and c_Th_ values (Fig. 8E) within the clusters but were within the range of values in the major clusters.

**Table 6:**
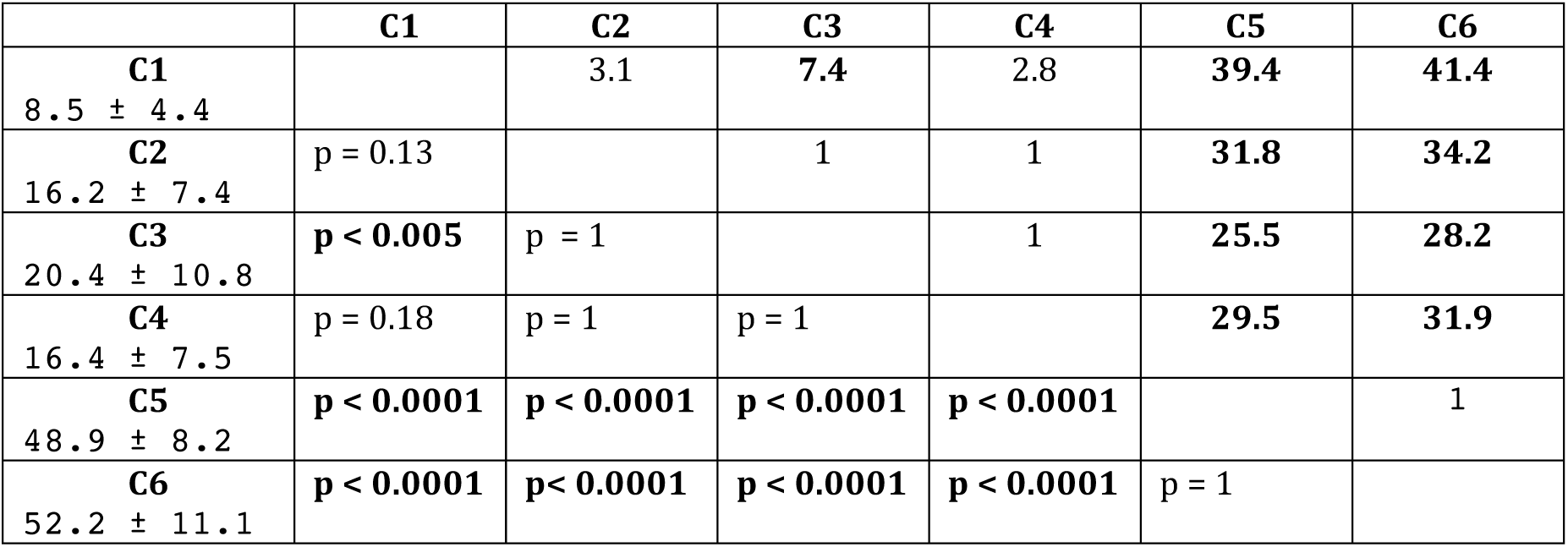
C50 Contrast: The values in the first column under the cluster number show the mean ± 1SD for each cluster. The p values were all determined as in Table 1.

#### Area tuning distributions

Most neurons in our sample summed visually evoked signals as stimulus size increased until a maximal area (Fig’s 1-3, E, J, O, T) and then maintained a plateau, with a small amount of suppression at the largest areas for a few neurons. Areal summation with weak or no surround suppression is characteristic of layer 6 neurons’ receptive fields (Gilbert 1977; Schiller et al, 1976a). The only cluster that showed a consistent pattern of suppression, albeit relatively weak (mean SI = 0.43 ± 0.22; Table 9), at the largest diameters was C1. The suppression observed in C1 neurons was significantly greater than in all the other clusters (Table 9).

**Table 7:** Peak Firing Rate: The values in the first column under the cluster number show the mean ± 1SD for each cluster. The p values were all determined using Kruskal Wallis non-parametric test. Other details as for table 1

|  | **C1** | **C2** | **C3** | **C4** | **C5** | **C6** |
| --- | --- | --- | --- | --- | --- | --- |
| **C1**<br>140 ± 43 |  | **12.3** | **13.2** | **12.3** | **10.5** | **9.9** |
| **C2**<br>52 ± 19 | **p < 0.0001** |  | 3.7 | **5.7** | **7.8** | **8.5** |
| **C3**<br>37 ± 16 | **p < 0.0001** | p = 1 |  | 2.5 | **4.8** | **5.6** |
| **C4**<br>29 ± 14 | **p < 0.0001** | p = 0.38 | p = 1 |  | 3.1 | 3.0 |
| **C5**<br>19 ± 9 | **p < 0.0001** | **p < 0.07** | p = 1 | p = 1 |  | 1.2 |
| **C6**<br>18 ± 7 | **p < 0.0001** | **p = 0.06** | p = 1 | p = 0.14 | p = 0.85 |  |

**Table 8:** Spontaneous Firing Rate: The values in the first column under the cluster number show the mean ± 1SD for each cluster. The p values were all determined using Kruskal Wallis non-parametric test. Other details as for table 1

|  | **C1** | **C2** | **C3** | **C4** | **C5** | **C6** |
| --- | --- | --- | --- | --- | --- | --- |
| **C1**<br>14.0 ± 7.3 |  | 3.6 | **15.1** | **13.3** | **10.5** | **10.1** |
| **C2**<br>9.0 ± 6.3 | p = 0.07 |  | **16.2** | **13.4** | **10.7** | **10.7** |
| **C3**<br>0.10 ± 0.20 | **p < 0.0001** | **p < 0.0001** |  | 2.6 | 2.4 | 2.1 |
| **C4**<br>0.51 ± 1.53 | **p < 0.0001** | **p < 0.0001** | p = 0.20 |  | 1.1 | 1.2 |
| **C5**<br>0.36 ± 0.82 | **p < 0.0001** | **p < 0.0001** | p = 0.24 | p = 0.88 |  | 1.2 |
| **C6**<br>0.09 ± 0.14 | **p < 0.0001** | **p < 0.0001** | p = 0.32 | p = 0.79 | p = 0.85 |  |

**Table 9:** Area Suppression Index: The values in the first column under the cluster number show the mean ± 1SD for each cluster. The p values were all determined using Student’s t-test. Other details as for table 1

|  | **C1** | **C2** | **C3** | **C4** | **C5** | **C6** |
| --- | --- | --- | --- | --- | --- | --- |
| **C1**<br>0.43 ± 0.22 |  | **6.6** | **7.7** | **8.3** | **5.0** | **4.8** |
| **C2**<br>0.19 ± 0.22 | **p < 0.005** |  | 1.8 | 1.9 | 1.0 | 1.1 |
| **C3**<br>0.14 ± 0.25 | **p < 0.025** | p = 0.47 |  | 1.1 | 1.5 | 1.5 |
| **C4**<br>0.13 ± 0.21 | **p < 0.0001** | p = 0.42 | p = 0.95 |  | 1.7 | 1.6 |
| **C5**<br>0.19 ± 0.23 | **p < 0.025** | p = 0.99 | p = 0.58 | p = 0.51 |  | 1.0 |
| **C6**<br>0.18 ± 0.24 | **p < 0.025** | p = 0.96 | p = 0.63 | p = 0.57 | p = 0.97 |  |

#### Response Rate and Spontaneous Rate

Seven functional tuning properties were extracted and z-transformed for the clustering but there were other tuning measures and functional response properties that were not used. A number of these are particularly notable because they tended to distribute according to cluster even though they were not included in the clustering analysis.

Consider maximum spike rate. Across the six main clusters there were mean maximum rates ranging from 140 spikes/sec down to 18 spikes/sec. The two main clusters accounting for the complex cell population (f1/f0 < 1) had clearly different ranges of peak firing rates. In the direction-selective complex cell cluster (C1) the mean rate was 140 ± 43 spikes/sec (Fig. 9A) which was significantly different from the mean rate of 52 ± 19 spikes/sec (Fig. 9C) in cluster C2 (p < 0.0001, Table 7).

**Figure 9:**
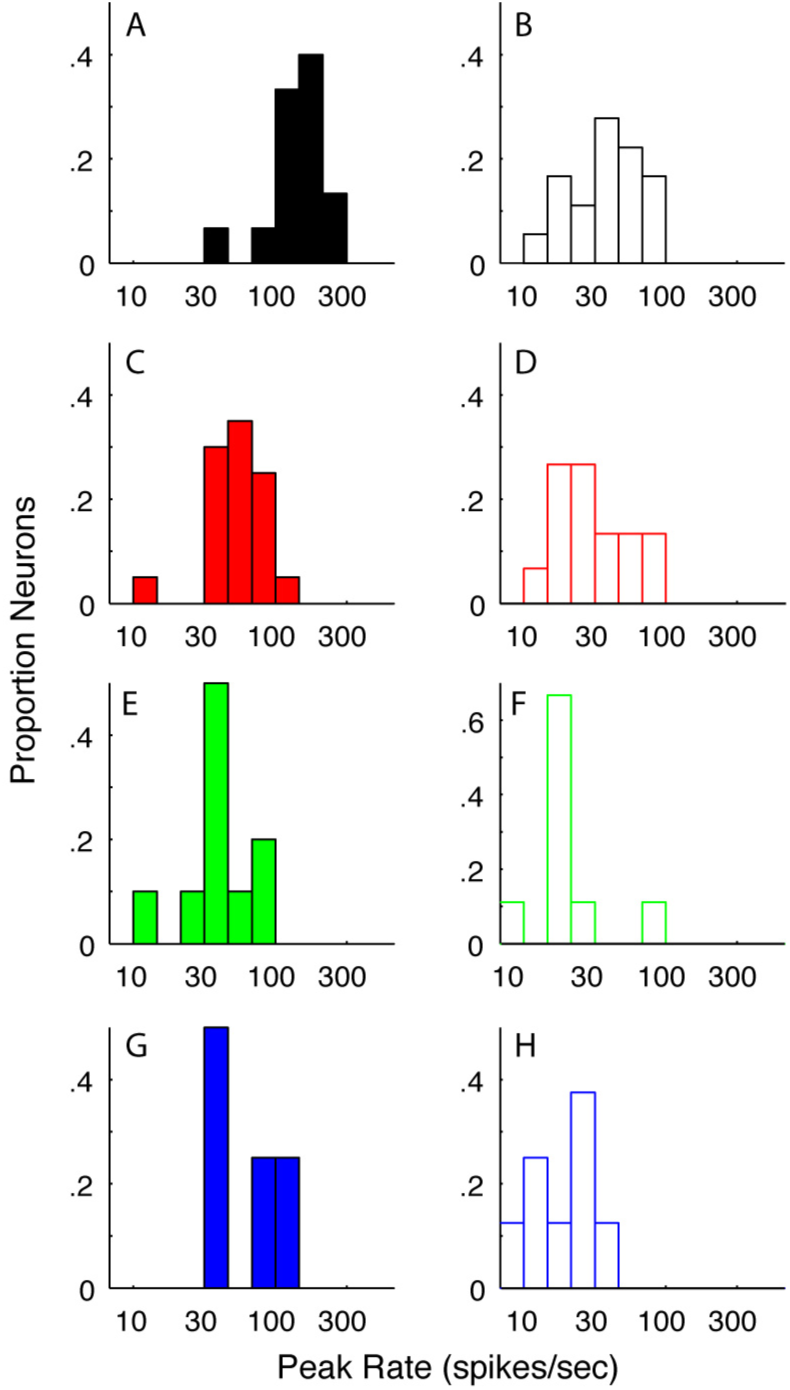
Distribution of maximum firing rates (see Methods) to optimal stimuli in each cluster. A, C, E, G:The distributions of peak firing rate of neurons in C1, C2, C6, C7 respectively all that had f1/f0 ratios < 1. Many neurons in C1 have high maximum firing rates. B, D, F, H:The distributions of peak firing rate of neurons in C3, C4, C5, C6 respectively, all had f1/f0 ratios > 1.

The remaining four major clusters that were predominantly simple cells. The larger two of these clusters (C3 and C4) had mean maximum spike rates 37 ± 16 and 29 ± 14 spikes/sec respectively which were not significantly different (p = 0.26,) while the two smaller clusters, C5 and C6, had mean peak rates of 19 ± 9 and 18 ± 7 spikes/sec respectively (cf. Table 7).

It is well established that spontaneous firing rate differs between simple and complex cell populations in the infragranular layers of macaque V1 (Schiller et al, 1976a, b; Ringach et al, 2002). We determined that this was the case for the layer 6 population (Fig. 10). The mean spontaneous rate for all neurons with f1/f0 ratios < 1 (complex cells) was 11.2 ± 8.2 spikes/sec whereas the mean rate for all neurons with f1/f0 ratios > 1 (simple cells) was 1.2 ± 3.7 spikes/sec. These distributions were significantly different (p < 0.0001). C1 had the highest spontaneous rate (14 ± 7.3 spikes/sec). C1’s spontaneous rate was not significantly different from the rate of C2 (9 ± 6.3 spikes/sec, Table 8). Of the overall population of neurons with f1/f0 ratios > 1, the four major simple cell clusters (C3-C6) all had similar ranges of f1/f0 and also had low mean spontaneous rates, all less than 1 spike/sec (Table 8) that did not differ between the clusters.

**Figure 10:**
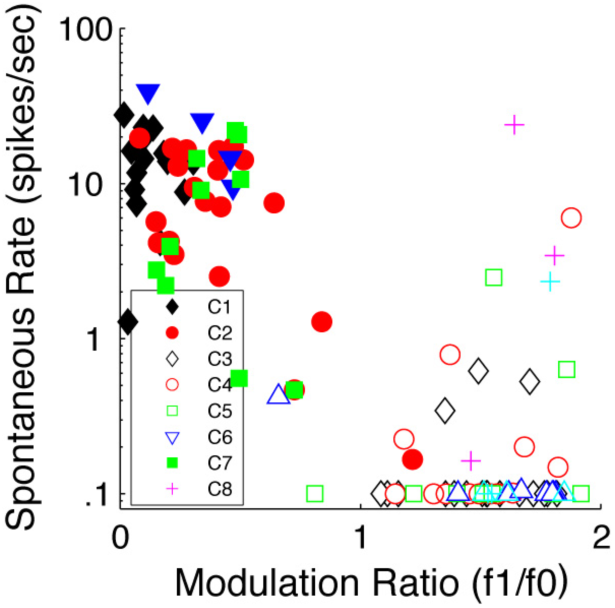
Distribution of spontaneous firing rates (y-axis) as a function of fl/f0 ratio (x-axis) for all neurons shown as clusters in Fig. 4. Neurons with spontaneous rate of 0.1 or less are shown as 0.1. There are two main groups, those with f1 /f0 < 1 have spontaneous rates greater than 1 and those with f1/f0 > 1 have low spontaneous rate. Clusters as in Fig 4.

### Relative Depth of Clusters within Layer 6

We determined whether the position of the recorded neurons within each cluster were distributed evenly across the depth of layer 6 or if there was any localization within the layer. This analysis was done by ordering the neurons in each cluster according to their relative depth within layer 6, and then comparing the resulting profile with the profile of the total population. The neurons in C1 were all found in the upper half of the layer as can be seen by comparing the black diamonds (C1) to the small blue circles (total population) in Figure 11A. In contrast, the neurons in C2 (Fig. 11A, red circles) were distributed through the depth of the layer. There did not appear to be any major aggregation of neurons in the other main clusters (C3 – C6) in the upper or lower regions of layer 6 (Fig. 11B).

**Figure 11:**
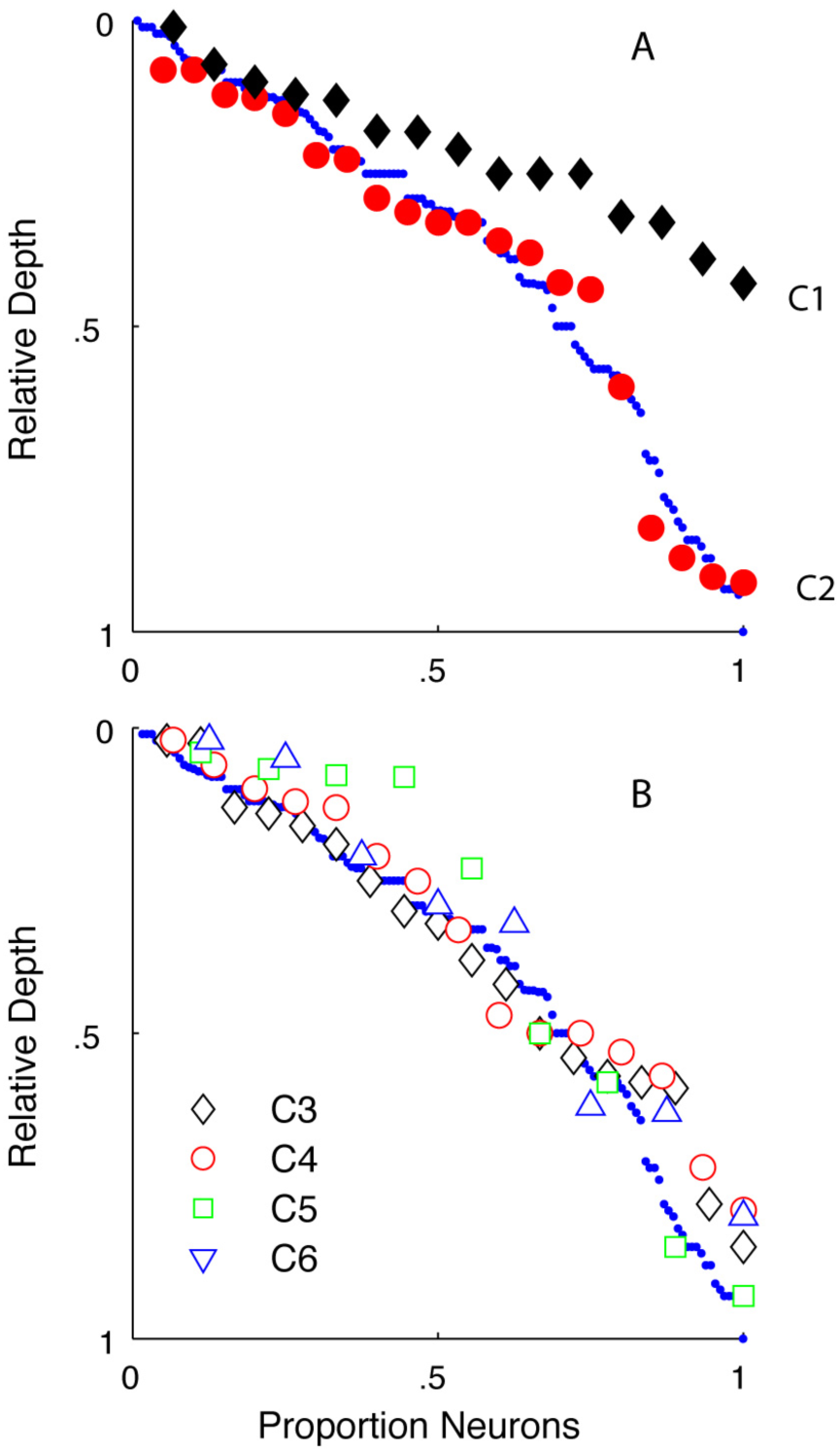
The relative depth of each neuron is shown in serial order from top of layer 6. Each cluster is shown by the same symbols as used in Fig. 4 and subsequent figures. A: The relative depth for C1 (black diamonds) shows that the neurons are found in the upper half of layer 6. Neurons in C2 (red circles) are distributed through the full depth of layer 6. The serial order of the total sample of 116 neurons is shown by the small blue circles. B:The four clusters with f1/f0 ratios > 1, C3 - C6 are distributed through the full depth of layer 6.

## Discussion

### Cluster analysis results

Of the seven tuning features used for cluster analysis, three measures were particularly useful for distinguishing the six major functional clusters in layer 6: modulation ratio (f1/f0), directional selectivity, and temporal frequency tuning.

The neurons in two of the six major clusters (C1 and C2) had f1/f0 ratios < 1 and would be classified as complex cells (Skottun et al, 1991). Neurons in the other four clusters (C3 – C6) had f1/f0 ratios > 1 and would be classified as simple cells.

A second feature defining the clusters was direction-selectivity. There were two major clusters – C1 and C3 – that had neurons that were strongly direction-selective (Fig. 5A,B, filled and unfilled black diamonds).

A third tuning measure, temporal frequency bandwidth, was also important in distinguishing C1 and C3 from the non-direction selective complex and simple cell clusters. C1 and C3 were bandpass in temporal frequency (Fig. 7A,B; Fig. 12, filled and unfilled black diamonds). Three of the remaining four major clusters (C2, C4 and C6) were either complex or simple (Fig. 4B; Fig. 12) but they were non-direction-selective (Fig. 5A,B; Fig. 12) and mostly low-pass in temporal frequency (Fig. 7A,B; Fig. 12).

**Figure 12:**
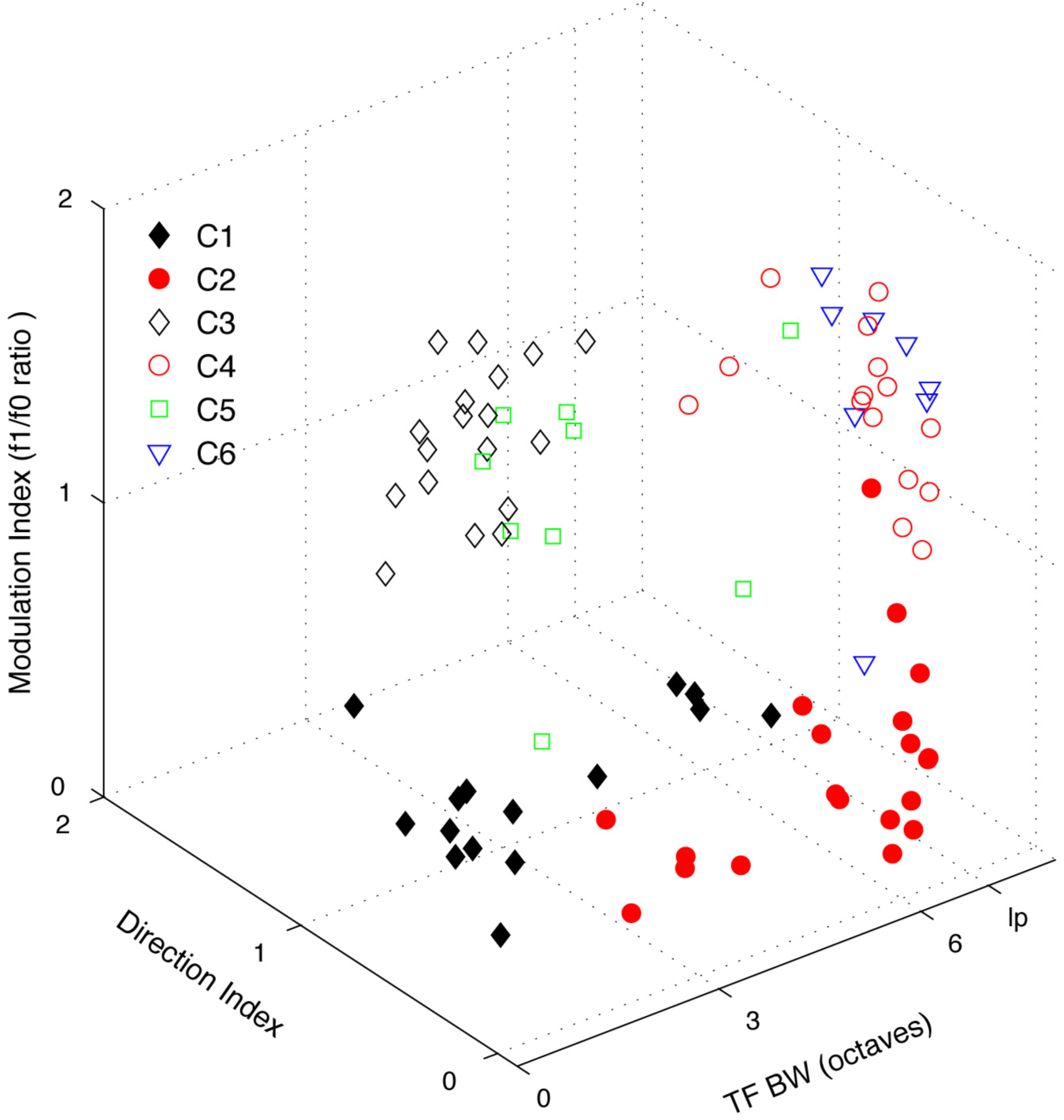
Distribution of the six largest clusters is shown with respect to three tuning parameters: temporal frequency bandwidth (x-axis), f1/f0 ratio (y-axis) and direction index (z-axis). Each cluster is indicated by the symbols described in Fig. 4.

### Tuning and response measures not used in the cluster analysis

Although we did not use firing rate as a parameter for clustering we showed in Results that visually-driven firing rate was much higher in C1 than in C2, and both clusters C1 and C2 had higher average firing rates than in the simple cell clusters C3-6. Similarly, spontaneous firing rate was significantly higher in C1 and C2 than in C3-6, and the separation between complex and simple cells within layer 6 (Fig. 10) was even more evident than the separation in the overall V1 population (Ringach et al, 2002). Furthermore, the distribution of cells in depth was different for C1 from that of all the other clusters (Fig. 11).

### Properties and possible functions of neurons in different layer 6 clusters

#### Cluster 1

The visually evoked mean rate was 140 spikes/sec for C1 neurons and they did not appear to adapt, much like the rates of PV inhibitory interneurons (Rudy and McBain, 2001). The non-adapting high spike rate of the PV inhibitory interneurons is attributed to the expression of the Kv3.1b potassium channel (Rudy and McBain, 2001). In rodents the Kv3.1b channel is not expressed in excitatory neurons in cortex. However, it is expressed in some excitatory neurons in primate cortex (Constantinople et al, 2009; Kelly et al, 2018), and it is clearly expressed in the large Meynert cells located near the layer 5/6 border in monkey visual cortex (Ichinohe et al, 2004; Constantinople et al, 2009; Kelly et al, 2018). It is tempting to speculate that some of the high firing rate neurons in C1 are large pyramidal neurons that express Kv3.1b. All neurons in C1 were located in the top half of the layer, layer 6a (Fig. 11).

Neurons in C1 were very direction-selective (Fig. 5C; Fig. 12). It has been established that most Meynert cells project to extrastriate visual area MT (Fries et al, 1985; Nhan and Callaway, 2012) which contains many direction-selective neurons (Zeki, 1974). Further, some of the tuning characteristics of the neurons in C1, such as strong directionality, low contrast threshold (Fig. 8A), and saturating contrast response functions (Fig. 1C, H), match those of the neurons described by Movshon and Newsome (1996) that were recorded in layer 6 of macaque V1 and that were activated antidromically by electrical stimulation of MT. The neurons in C1 showed some response-attenuation at large stimulus window diameters (Fig. 1E,J) indicative of eCRF suppression. Their eCRF suppression may be inherited from magnocellular LGN neurons because M-pathway retinal ganglion cells (presumptive parasol cells) show moderate eCRF suppression (Solomon et al, 2006). If some of the C1 population are also providing input to MT it is consistent with the findings that the functional characteristics of the input to MT neurons are derived mainly via V1, V2 and V3 through the M-pathway (Gegenfurtner et al, 1994).

#### Cluster 2

Neurons in C2 were also complex cells (f1/f0 ratio < 1) but were not direction-selective (Fig. 1K, P). However, all C2 neurons were orientation- and spatial-frequency-selective (Fig. 1L,Q). The mean spike rate of the neurons in C2 was significantly lower than the mean rate in C1 (Table 7). As a cluster they had higher average cTh’s and c50’s than C1 (Fig. 8; Tables 5 and 6), but the differences were not significant. Sixty-five percent of the neurons in C2 were lowpass in temporal frequency (13/20, Fig. 1N) and the majority showed little eCRF suppression: the mean SI was 0.2, but there were two neurons that had SI > 0.7. In distinction to the neurons in C1, this group of neurons would provide signals about stationary or slowly moving borders.

#### Cluster 3

All the neurons in C3 were simple cells (f1/f0 ratio > 1, mean = 1.5) that were direction-selective, orientation- and SF-selective and often strongly bandpass in TF (Fig. 2A-J). The average orientation bandwidth was 16 deg (Fig. 5D; Table 1), showing the narrowest average tuning of all the clusters. The neurons in C3 also had narrow TF bandwidths (mean 2.3 octaves, Table 4) but a range of optimal TF’s (Fig. 7B). The combination of narrow tuning for TF but a range of optimal TF’s and a range of optimal SF’s (Fig. 6B) suggests that this population of simple cells may span a selective range of velocity preferences. This cluster of neurons would signal the sign and direction of movement of objects and edges over all but the lowest contrast range.

#### Cluster 4

The neurons in C4 were distinguished from those of C3 by their lack of direction selectivity (Fig. 2K, L, N, P, Q, S; Fig. 5D) and their lowpass temporal frequency tuning (Fig. 7B; Table 4). The cluster had a similar range of contrast threshold and c50 to C3. The average peak firing rate of neurons in C4 was not significantly different from the rate in C3 (Table 7). The neurons in C4 were significantly more sensitive to contrast than those in C5 and C6 (Tables 5, 6). This cluster of neurons, because they are simple cells and are lowpass in temporal frequency, are the only cluster that would signal the sign and location of oriented edges in an image across a range of sizes at low and intermediate contrasts in a stationary achromatic image.

#### Clusters 5 and 6

Most neurons in C5 and C6 were characterized by relatively narrow orientation and spatial frequency bandwidth combined with low sensitivity to achromatic contrast. It has been a longstanding mystery how the visual system maintains a high level of contrast discrimination for achromatic stimuli over a wide range of base contrasts (Legge, 1981; Barlow et al 1987; Chirimuuta and Tolhurst, 2005a, b). The parcellation of neurons into clusters with different dynamic ranges for different contrast ranges suggests a solution to this long-standing puzzle of contrast discrimination and identification (Chirimuuta and Tolhurst, 2005a, b). Clusters C5 and C6 were distinguished from each other by their temporal frequency selectivity. Almost all neurons in C5 were bandpass for TF (Fig. 7D, green squares) whereas those in C6 were lowpass for TF (Fig. 7D, blue triangles). Both clusters were distributed through the depth of layer 6 (Fig. 11B).

#### Minor Clusters

Some of the smaller clusters had neurons that were relatively untuned for orientation and lowpass in spatial frequency (Fig. 3K, L, P, Q; Fig, 5E, G; Fig. 6C, E). These are properties that have been attributed to inhibitory interneurons (Nowak et al, 2008) but anatomical identification along with functional characterization is necessary to show that these neurons are inhibitory.

### Implications of clusters

If the functional subpopulations described in the current study are related to the structurally defined subclasses in layer 6 (Wiser and Callaway, 1996; Hasse et al, 2018) then there will be separate target regions for the axons of these neurons. Many layer 6 neurons project intracortically (Wiser and Callaway, 1996) and some have axons preferentially terminating in layer 4cα – the major M-pathway recipient sublayer in V1 – while others that preferentially arborize in layer 4cβ and layer 4a – the principal P-pathway recipient zones of layer 4. It is not known how these anatomical subclasses map onto our functionally defined clusters. Around 10 - 15% of layer 6 neurons provide feedback projections to the LGN (Fitzpatrick et al, 1994) and these have some functional attributes (Briggs and Usrey, 2009; Hasse and Briggs 2017) that align mainly with two of the clusters (C2 and C6) we have identified. Another structurally defined group of neurons project to extrastriate visual area MT (Lund et al, 1975; Fries et al, 1985; Nhan and Callaway, 2012) and Movshon and Newsome (1996) showed that some layer 6 neurons projecting to MT have functional properties similar to those of C1. Future studies combining functional and anatomical characterization may reveal whether these functional clusters indeed correspond to particular anatomical classes.

## Acknowledgements

This work was supported by NIH grants EY1472, EY8300, EY15549 to RMS and MJH and core grant P30 EY13079 and training grant T32 EY7136.

